# Within-species phylogenetic relatedness of a common mycorrhizal fungus affects evenness in plant communities through effects on dominant species

**DOI:** 10.1101/328708

**Authors:** Romain Savary, Lucas Villard, Ian R. Sanders

**Affiliations:** Department of Ecology and Evolution, Biophore Building, University of Lausanne, 1015 Lausanne, Switzerland.

**Keywords:** AMF, *Rhizophagus irregularis*, plant community, plant-plant interactions, phylosignal, evenness

## Abstract

Arbuscular mycorrhizal fungi (AMF) have been shown to influence plant community structure and diversity. Studies based on single plant - single AMF isolate experiments show that within AMF species variation leads to large differential growth responses of different plant species. Because of these differential effects, genetic differences among isolates of an AMF species could potentially have strong effects on the structure of plant communities.

We tested the hypothesis that within species variation in the AMF *R. irregularis* significantly affects plant community structure and plant co-existence. We took advantage of a recent genetic characterization of several isolates using double-digest restriction-site associated DNA sequencing (ddRADseq). This allowed us to test not only for the impact of within AMF species variation on plant community structure but also for the role of the *R. irregularis* phylogeny on plant community metrics. Nine isolates of *R. irregularis*, belonging to three different genetic groups (Gp1, Gp3 and Gp4), were used as either single inoculum or as mixed diversity inoculum. Plants in a mesocosm representing common species that naturally co-exist in European grasslands were inoculated with the different AMF treatments.

We found that within-species differences in *R. irregularis* did not strongly influence the performance of individual plants or the structure of the overall plant community. However, the evenness of the plant community was affected by the phylogeny of the fungal isolates, where more closely-related AMF isolates were more likely to affect plant community evenness in a similar way compared to more genetically distant isolates.

This study underlines the effect of within AMF species variability on plant community structure. While differential effects of the AMF isolates were not strong, a single AMF species had enough functional variability to change the equilibrium of a plant community in a way that is associated with the evolutionary history of the fungus.

## Introduction

Soil microorganisms that influence plant-plant interactions play a central role in terrestrial ecosystems [1]. This is particularly true for arbuscular mycorrhizal fungi (AMF; phylum Glomeromycota), which are considered the commonest of plant root symbionts, due to their unique capacity to form endosymbioses and to exchange nutrients with 74% of land plants [2]. During the last decades, various beneficial effects of these fungi on different plant species were reported such as an increase in plant growth and plant nutrient acquisition, [3], greater resistance to pathogens [4] and herbivores [5], and increasing tolerance to drought, high salinity and pollutants [6–8]. In addition to direct effects on plant physiology, AMF have also been shown to alter competitive interactions between plants [3]. Consequently, this impacts common metrics of plant community structure, such as community richness, community evenness (i.e the relative abundance of community members) and it affects also the community productivity. For example, removing AMF from nutrient-poor tallgrass prairies, where the dominant plant is highly mycotrophic, had a tendency to favor facultative mycotrophic plants, thus increasing total community evenness and richness without increasing total productivity [9]. Conversely, in grasslands, dominated by grass species that derive little benefit from the association, AMF have been observed to favour the productivity of subordinate forbs [3], thus increasing community evenness and richness [10]. Other studies also demonstrated that AMF could promote or limit community productivity depending on the AMF taxon involved, regardless of plant community species richness [11].

In experiments studying the effect of an AMF species inoculated on a single plant species, the range of growth responses of plants was found to vary greatly depending on the combination of plant species and AMF species tested; spanning a range of response from highly positive to negative [12]. One possible explanation for this effect was that AMF species were not equivalent in their functional characteristics [13] and, thus, impacted the outcome of a specific AMF-plant interaction. Later on, Powell et al. (2009) [14] showed that some of these differences were correlated with AMF phylogeny. Indeed, they found that some fungal quantitative traits of members of the Glomeromycota phylum appear to be phylogenetically conserved at the family level. Consequently, this created a similar conservatism towards plant response. Similarly, it was shown that the plant response varied greatly while in symbiosis with different genotypes of a single AMF species [15–19]. Depending on the species tested, the amplitude of variation in plant response due to within-AMF species differences was similar, or even higher, than the variation in effects among AMF species. These observations lead Sanders and Rodriguez (2016) [19] to suggest that there is an inconsistency between the level of AMF phylogenetic resolution used by experimental ecologists and the level of functionality in the AMF phylogeny. Indeed, most ecological studies to date used one representative isolate for each species [1, 20–21], making the assumption that the within-AMF species genetic diversity and its plant effect was homogenous across any isolate of that species. In the light of previous results [15–16, 18] this assumption would seem to be compromised. From the observed effects of high variation in a plants response to different genotypes of a single AMF species, we could expect that different *R. irregularis* genotypes might differentially affect plant-plant interactions in a plant community. Moreover, according to the findings of Powell et al. (2009) [14], we could also expect that intra-specific variation in fungal traits, plant response and community response might be explained by phylogenetic conservatism. Testing intraspecific phylogenetic effects on plant communities were previously difficult because of the lack of accurate data on intraspecific genomic differences in AMF.

According to recent estimates, the Glomeromycota phylum is composed of 300 to 1600 AMF species [2] and the vast majority of ecosystems harbour an assembly of several AMF species. Studies that manipulated AMF species richness in a plant community have shown that there was a positive relationship between AMF diversity and community productivity [3]. This effect could be the result of a functional complementarity of co-occuring taxa, since Maherali and Klironomos (2007) [22] have shown that more phylogenetically diverse AMF communities were more stable and resulted in greater plant biomass compared to the effects of AMF communities composed of taxa that were phylogentically similar. In the putative framework of a phylogenetic conservatism of AMF traits and plant responses, one could expect that AMF communities that are phylogenetically diverse at the intraspecific level will positively impact the different community metrics, as it has been suggested that a mix of different genotypes could affect ecosystem function as much as a mix of species [22].

In this study, we took advantage of a recent characterization of intraspecific genetic variability in one of the commonest AMF species *Rhizophagus irregularis*, using double-digest restriction-site associated DNA sequencing (ddRADseq) [23]. We also chose to use this AMF species because of the previously documented variability in quantitative traits and effects on plant growth [17]. We tested the importance of intraspecific diversity of *R. irregularis* on different plant species co-existing within a plant community and on characteristics of plant communities. We sought to determine if genetic relatedness among isolates of *R. irregularis* was correlated to the putative effects on single plant species within a plant community and on characteristics of mesocosm plant communities. We also wanted to test if increasing phylogenetic diversity in a community of *R. irregularis* could positively affect plant community productivity and evenness.

We built artificial mesocosms with a fixed plant community composed of six plant species, that are typical of central European calcareous grasslands. We chose nine *R. irregularis* isolates equally distributed into three genetic groups describing a large part of the known *R. irregularis* intraspecific genetic variation [23]. The genetic groups are named Gp1, Gp3 and Gp4 [23]. Both Gp3 and Gp4 naturally co-exist in several locations and all three genetic groups occur in central Europe [23]. All isolates were cultured for several generations in an *in vitro* culture system in order to remove potential environmental effects from their place of origin that could otherwise be confounded with genetic effects. These isolates were either inoculated singly into the mesocosms or as a combination of isolates. This design allowed us to test three main questions: i) Do different isolates of *R. irregularis*, or combinations of the isolates, differentially colonize plants and differentially affect plant responsiveness, community structure and productivity? ii) Do individual plant species of a community respond more similarly to more closely genetically related AMF isolates compared to more distant ones? iii) Is the mycorrhizal responsiveness and structure of a plant community more similar in response to more closely genetically related AMF isolates compared to more distant isolates?

## Material & Methods

### Soil, fungal treatments, plant material and experimental design

A natural clay soil collected from a calcareous grassland at the University of Lausanne, Lausanne, Switzerland (46°31’32.0016" N, 006°34’46.6068" E) was used for this experiment. The soil was sieved through a 1 cm mesh, then mixed with quartz sand in a ratio 2:3. This mixture was then steam-sterilized at 105°C for three consecutive sessions of 20 min. Round 4.6 litre pots (30 cm diameter) were filled with the sterilized soil:sand mixture.

Nine *R. irregularis* isolates were chosen in three different genetic groups (Gp1, Gp3 and Gp4), representing the major part of the diversity described in this species [23]. These were: Gp1 - 5B, ESQLS69, LPA54; Gp3 - A4, DAOM229457, DAOM240409; Gp4 - A1, DAOM197198-CZ, DAOM240159 (Fig. 1a). The delineation of this variation was based on previous ddRADseq data generated from DNA of 59 isolates of *R. irregularis* isolated from several geographical locations and comprising 6888 sites in the genome where single nucleotide polymorphisms (SNPs) occurred. Each of the 9 isolates was first grown in an identical *in vitro* environment for 5 months [24]. In order to create a natural soil inoculum for each isolate [25], we inoculated *P. lanceolata* with 500 *in vitro*-produced spores in ten 0.5 litre pots in a sterile autoclaved Oil-Dri^®^ granular clay (Oil Dri corporation of America) with quartz sand in a 3:2 ratio. Ten *P. lanceolata* were mock inoculated with water in order to produce a substrate representing a mock-inoculated treatment. The *P. lanceolata* plants were grown for five months and then the soil of the ten pots per isolate were mixed to create the 9 inoculum stocks and substrate for the mock-inoculated treatment.

**Figure 1.**
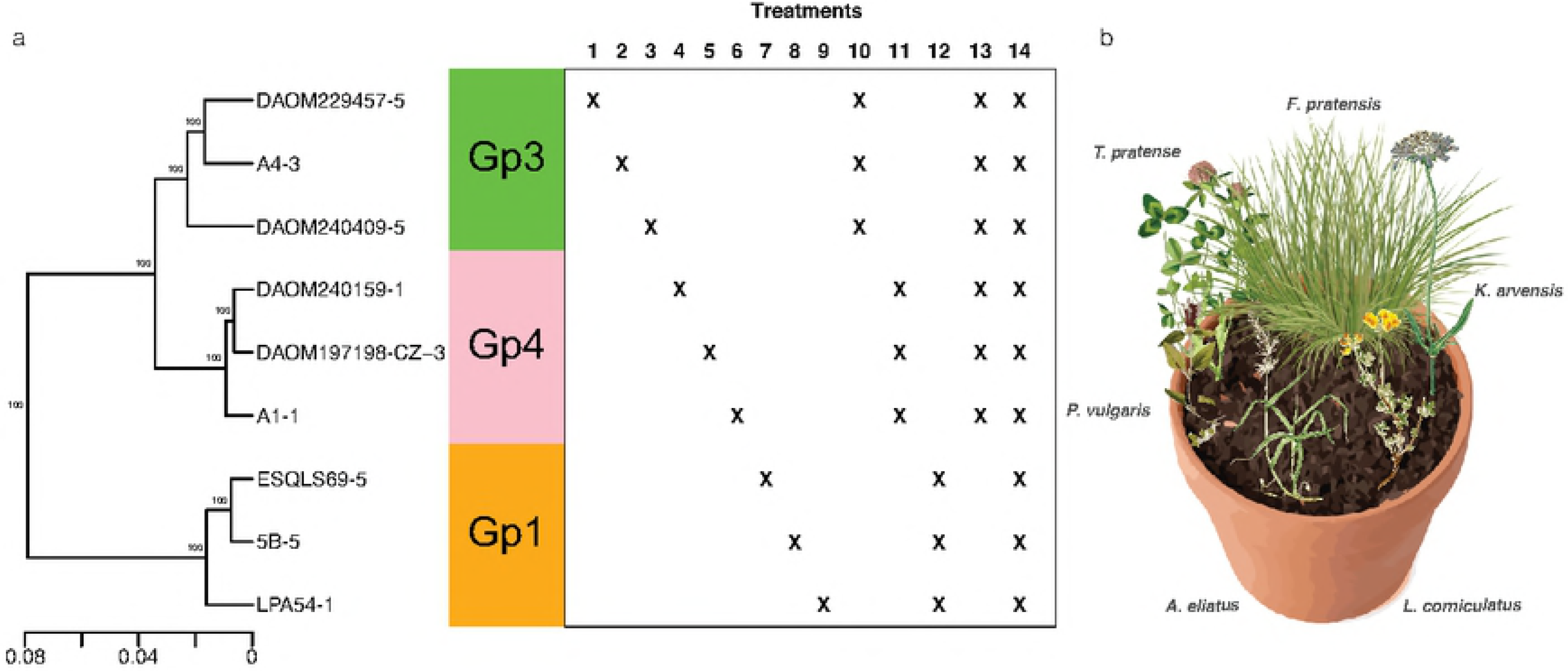
Study design. **(a)** Phylogeny of *R. irregularis* isolates from three genetic groups used as treatments either as single inoculants (treatments 1-9) or as mixed inoculant (treatments 10-14). **(b)** Representation of one mesocosm with the six plants.

In order to construct simple artificial European calcareous grassland community in the greenhouse, we chose six plant species commonly found in this type of phytosociological association (classified as *Arrhenatherion* [26]). The following plant species were chosen to be equally partitioned across the different functional groups [27]. Thus, we chose two legumes, *Trifolium pratense* and *Lotus corniculatus*, two grasses, *Arrhenatherum elatius* and *Festuca pratensis* and two forbs *Prunella vulgaris* and *Knautia arvensis*. All these species are known to form the arbuscular mycorrhizal symbiosis [28]. Seeds of these plants were obtained from UFA Samen (Switzerland) and germinated in trays on the same substrate used for the experiment. After two weeks of growth, one individual of each species was planted in a circle in each pot (Fig. 1b). The position was randomly chosen except that plants of the same functional group were never planted next to each other but always at the most distant location. Thus, *Arrhenatherum elatius* was always the most distant plant from *Festuca pratensis, Trifolium pratense* was always the most distant plant from *Lotus corniculatus* and *Prunella vulgaris* was always the most distant plant from *Knautia arvensis* (Fig. 1b).

Fourteen mycorrhizal treatments and one non-mycorrhizal (NM) treatment were applied to the plant communities and were replicated ten times (Fig. 1a). We inoculated each pot of the 9 single inoculation treatments (treatments 1-9) with 100g of inoculum. Treatments with a co-inoculation of three isolates (Gp1, Gp3 and Gp4; treatments 10-12) received 33.3g of inoculum of each isolate. In the case of Gp3+Gp4 (treatment 13) we used 16.6g of inoculum of each of the six isolates. Finally in the co-inoculation treatment with all isolates (treatment 14), 11.1g of inoculum of each isolate was added per pot. The fifteen treatments were randomly arranged on a table in the greenhouse. This procedure was repeated 10 times for the 10 replicates separated on 10 tables. Tables were regularly randomized to avoid microclimate effects. The greenhouse conditions were constant at, 24°C, 60% RH and 12h of daylight. The communities were grown for 3.5 months from May 4th 2015 to the August 19th 2015.

### Harvest, measurements and statistical analysis

After 105 days of growth, the shoots of each plant in the community were harvested and dried at 70°C for three days and weighed to obtain the above ground dry mass (ADM) for each of the 900 plants. A small part of the roots of each plant was collected and frozen at −20°C for later measurement of AMF colonization by staining. Roots were stained and AMF colonization was measured (see supplementary note S1). The roots from each mesocosm of all the six plants were washed dried together and weighed, thus, giving the total community root dry mass (RDM). The sum of all plant ADM from one pot plus the RDM of this pot resulted in total dry mass (TDM) of the community. Inflorescence number was counted at 78 and 105 days of growth. In order to have a standardized measure of mycorrhizal effects on the different plant species, plant individual responsiveness was calculated following Gange and Ayres (1999) [29] based on the mean ADM for each plant species in the non-mycorrhizal treatment. We used the data collected for each individual in a mesocosm to build the metrics of community structure. TDM was considered as the community productivity. Mean community responsiveness and mean AMF colonization were obtained by averaging the individual responsiveness across plant species and AMF colonization across plant species, respectively. Community evenness was measured with Pielou’s evenness index [30] using the diversity function in vegan 2.3-3 [31]. This was calculated by dividing the Shannon index of ADM by the log of the number of individuals in the community. Evenness represents a value of equality (1) or inequality (0) of biomass partitioning among species within a mesocosm. The AMF colonization evenness was calculated in the same way in order to assess equality or inequality of AMF colonization among the six plant species within a mesocosm. This could be considered as a proxy for AMF preferences within a mesocosm. Significant correlations between mesocosm variables, as well as the relationships between single plant data and mesocosm averages, were assessed using Pearson’s product moment and polynomial regressions.

The effect of plant species and mycorrhizal treatments on AMF colonization, plant responsiveness and number of flowers produced were analyzed using linear mixed-models (lmer) with the pot as a random factor. In order to test for significant differences an analysis of variance was used on the models. Significant pairwise comparisons were assessed using the difflsmeans function. Significant differences among AMF treatments towards plant community metrics were assessed using a one-way ANOVA and a Tukey HSD post-hoc test. All statistical analyses were performed in R, version 3.4.4.

### R. irregularis *phylogeny and phylogenetic signal*

All nine isolates used in this study were previously sequenced with ddRAD-seq with a minimum of 3 replicates [23]. The raw sequence reads were retrieved from the NCBI bioproject (accession number PRJNA326895) and were trimmed and analysed following the workflow of Wyss et al., (2016) [32] and Savary et al. (2017) [23]. Genetic distance matrices among the nine isolates were calculated based on the scalar distance method of Wyss et al., 2016 [32]. For this calculation, data used were taken from the replicate with the deepest sequencing of each of the nine isolates and with the highest number of markers available.

The package phylosignal [33] was used to test phylogenetic signals between the phylogeny of the 9 isolates and i) AMF colonization and mycorrhizal responsiveness across the 6 plant species and ii) the mean AMF colonization, mean mycorrhizal responsiveness, evenness, total productivity and mean flower production per mesocosm, 78 and 105 days after inoculation. Five phylogenetic signal indicators were calculated on variables of each of the plant species and on community metrics of the mesocosms. These were Moran’s I index [34,35], Abouheif’s Cmean index [36], Blomberg’s K and K* [37] and Pagel’s λ [38]. These were then tested against the null hypothesis of a random trait with no significant signal [33]. Plant and mesocosm variables showing significant phylogenetic signal towards AMF isolates were kept for following analyses on co-inoculation treatments. AMF community phylogenetic diversity for treatments involving more than one AMF taxa was calculated using Faith’s PD [39] and relationships between the retained variables and phylogenetic diversity were assessed using quantile polynomial regression on the median, 20th and 80th quantiles of the response variable.

## Results

### Effect of fungal treatments on individual plant species and on plant communities

Overall, plants in the fourteen treatments exhibited high levels of AMF colonization in all six plants with a global mean of 67.6% ± 19.7% (SD). Fifty-seven plants in the NM treatment exhibited no AMF colonization. Three plants were measured with a very small amount of AMF colonization (less than 5%). These were considered as a contaminant. Colonization of the roots by the fungi was significantly affected by the different AMF treatments as well as by plant species identity. However, the interaction between these two variables was not significant (Table 1a). Isolates A4 (Gp3) and ESQLS69 (Gp1) were significantly the lowest colonizers and the three isolates of Gp4: A1, DAOM240159 and DAOM197198-CZ were the highest colonizers (Fig. 2a). *R. irregularis*, independent of isolate identity, colonized a significantly greater proportion of the roots of *K. arvensis, P. vulgaris* and *T. pratense* than the dominant plant, *F. pratensis* (Fig. 2b).

**Table 1.** **(a)** Results of ANOVA on the mixed-models including the pot as a random factor for the variable measured on individuals plants: AMF colonization, mycorrhizal responsiveness and flower production at 78 and 105 days per plant; **(b)** Results of ANOVA for the seven mesocosme variables: above-ground dry mass evenness, root dry mass, above-ground dry mass, total productivity, flower production at 78 and 105 days, and AMF colonization evenness.

| Source of Variation | df | MS | F |
| --- | --- | --- | --- |
| **a. Variables measured on individuals plants** |  |  |  |

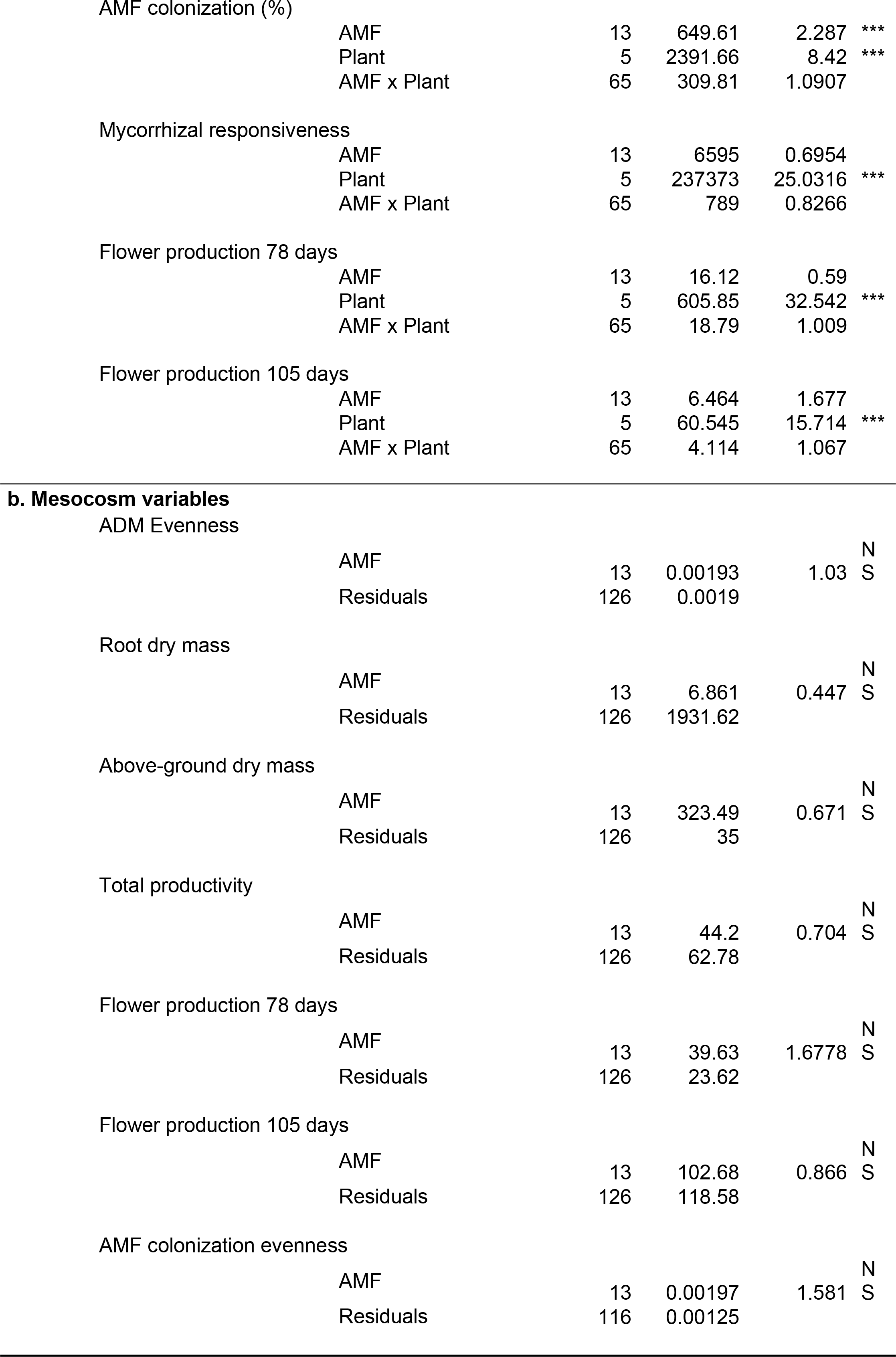

**Figure 2.**
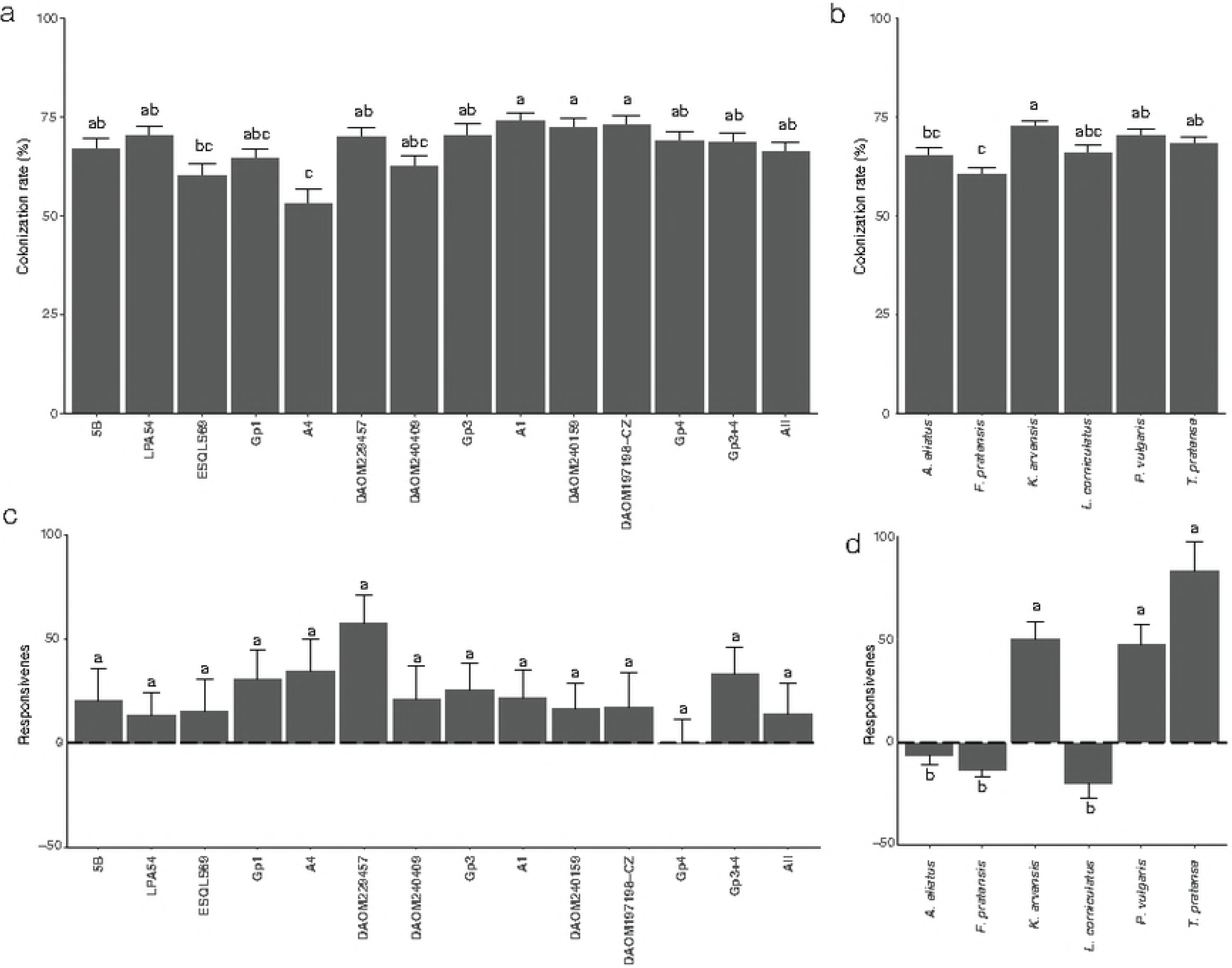
AMF colonization and responsiveness in the different treatments and different plant species. **(a)** Mean and the standard error of AMF colonization (% root length) according in the 14 treatments, and **(b)** in the six plants species. **(c)** Mean and the standard error of the responsiveness of each plant in the different treatments and **(d)** in the six different plant species of the community.

Large and significant differences in mycorrhizal responsiveness occurred among the different plant species but was not significantly influenced by the different AMF isolates (Table 1a). Mycorrhizal responsiveness of the plant community was positive (23.13 ± 107.82 SD) in the majority of fungal treatments (Fig. 2c). Mycorrhizal responsiveness of the grasses *A. eliatus* and *F. pratensis*, and *L. corniculatus* was negative and differed significantly from the positive mycorrhizal responsiveness of *K. arvensis, P. vulgaris* and *T. pratense* (Fig. 2d). *F. pratensis* responsiveness decreased with increasing AMF colonization (cor= −0.21, p= 0.013; Fig. S1). In contrast, mycorrhizal responsiveness of *P. vulgaris* and *T. pratense* increased with increasing AMF colonization (cor= 0.30, p<0.001, cor= 0.25, p= 0.004, Fig. S1). There was no significant correlation between mycorrhizal responsiveness and AMF colonization in the remaining plant species.

The dry mass production of the mesocosms either as RDM, ADM or as total productivity (TDM), was not significantly affected by any of the AMF treatments (Fig. S2a and S2b, Table 1b). Plant community evenness and AMF colonization evenness were not significantly different across all treatments (Fig. S2c; Table 1b). However, a number of interesting correlations were observed between variables. Mean AMF colonization per mesocosm was significantly and positively correlated with plant community evenness (cor=0.256, p=0.0023, Fig. S3), as was mean AMF colonization per mesocosm with AMF colonization evenness (cor=0.724, p<0.001, Fig. S4). Mean mesocosm responsiveness was significantly and positively correlated with mean AMF colonization (cor=0.300, p<0.001, Fig. 3a) and total productivity (cor=0.481, p<0.001, Fig. 3b). Plant community evenness was significantly and positively correlated with mean mycorrhizal responsiveness (cor=0.752, p<0.001, Fig. 3c). The increase in plant community evenness was associated with a strong significant decrease in the relative contribution of *F. pratensis* to the total above-ground mesocosm productivity and a significant increase in the relative contribution of all the other plants (Table S1; Fig. 3d).

**Figure 3.**
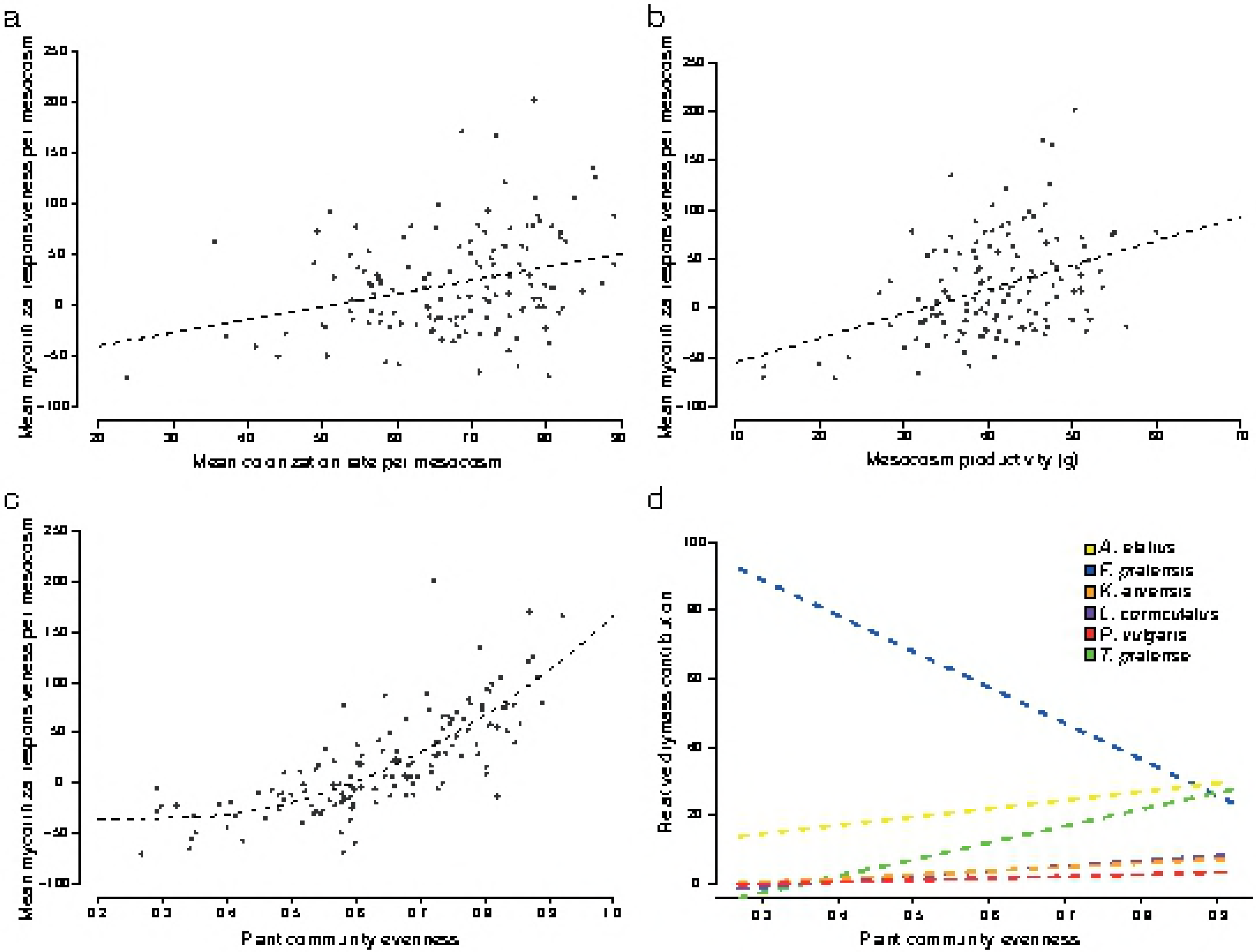
Community metrics and relative dry mass contribution of each plant to the community. **(a)** Correlation between the mean mycorrhizal responsiveness and the mean level of AMF colonization (cor=0.300, p<0.001) **(b)** Correlation between the mean plant responsiveness and the mesocosm productivity (TDM, cor=0.481, p<0.001) **(c)** Quadratic relation between the mean plant responsiveness and the plant community evenness per mesocom (cor=0.752, p<0.001). **(d)** Relative dry mass contribution of each plant species to the mesocosm according to the estimator of plant community evenness. All models were significant (p<0.001)

### Phylogeny of the nine isolates

Out of the ddRAD-seq data [23] from the nine isolates used in this study, we retrieved the data from the biological replicate of each isolate that had been sequenced the deepest and with the highest SNP calling quality. We were able to retrieve 60668 shared sequence positions, which contained 16311 SNPs, 1137 insertions, 1453 deletions and 11 MNPs. The much higher number of variable positions, in comparison to the study of Savary et al. (2017) [23], was due to the low number of isolates and replicates used. This enabled us to retrieve shared positions among all isolates that would not be shared among all isolates in a larger data set. This variation was used to build 100 genetic distance matrices from 5000 randomly chosen sites, following the method described in Wyss et al., (2016) [32]. All nodes showed a support value of 100, clearly separating the nine isolates into three main groups (Fig. 1a). This tree was then used for phylogenetic signal analyses.

### Testing for phylogenetic signals with plant species data and plant community metrics

There was a significant within-species AMF phylogenetic signal in AMF colonization of *F. pratensis* in two out of the five tests (Fig. 4a), with a globally highest colonization by Gp4 isolates. Similarly, there was a significant within-species AMF phylogenetic signal in the mycorrhizal responsiveness of *F. pratensis* (Fig. 4b) in three tests out of the five. There was also a significant within-species AMF phylogenetic signal in the mycorrhizal responsiveness of *K. arvensis* for one test out of five (Fig 4b).

**Figure 4.**
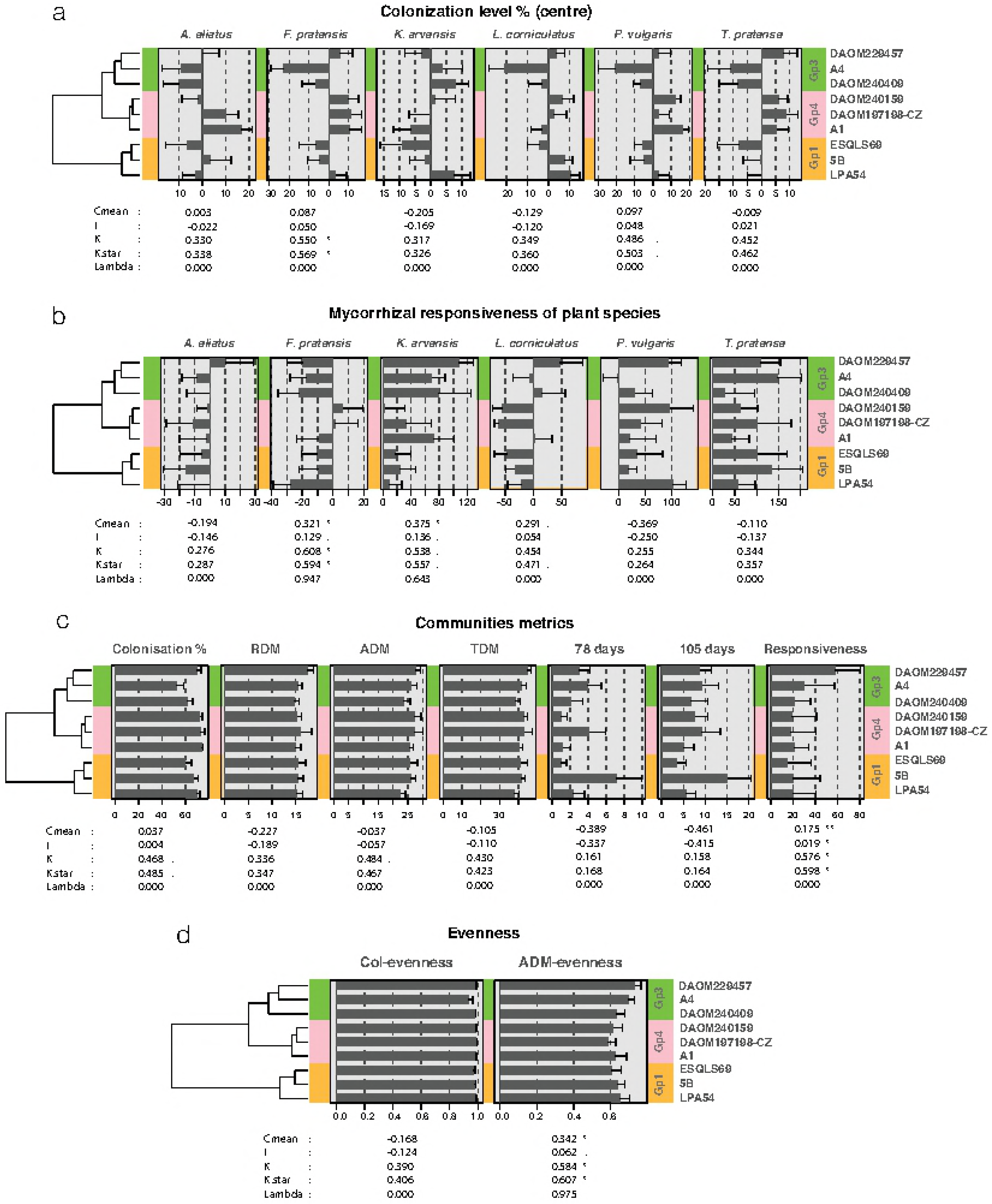
Individual plant species and mesocosm phylogenetic signals. Each of the phylogenetic signal plots are composed of the phylogeny of the nine isolates from the three genetic groups. The value of the different tests and their significance (. < 0.1, * < 0.05, ** < 0.01) are indicated under each set of bar plots. The phylosignals were calculated for each of the six plants independently for **(a)** AMF colonization (%), that was centred on the mean colonization per plant, and **(b)** for the responsiveness (not centered). Mesocosm phylosignals are presented **(c)** for mean colonization, RDM, ADM, TDM, flower production after 78 and 105 days and mean mesocosm responsiveness and **(d)** for mesocosm colonization repartition (Col-evenness) and plant community evenness (ADM-evenness).

At the community level, the overall mean AMF colonization, RDM, ADM, TDM and flower production did not reveal a significant within-species AMF phylogenetic signal (Fig. 4c). In contrast, there was a significant within-species AMF phylogenetic signal in mean mycorrhizal responsiveness of the community in four out of five tests (Fig. 4c). There was no detectable within-species AMF phylogenetic signal in AMF colonization evenness (Col-evenness) (Fig. 4d). There was a significant within-species AMF phylogenetic signal in terms of community evenness (ADM-evenness) in three tests out of five (Fig 4d). Phylogenetic diversity of the AMF community, calculated with Faith’s index, showed a significant quadratic relationship with plant community evenness (y=0.023x^−0.088^ + 0.195, p=0.0475, Fig. 5), showing the highest community evenness values at an intermediate level of phylogenetic diversity.

**Figure 5.**
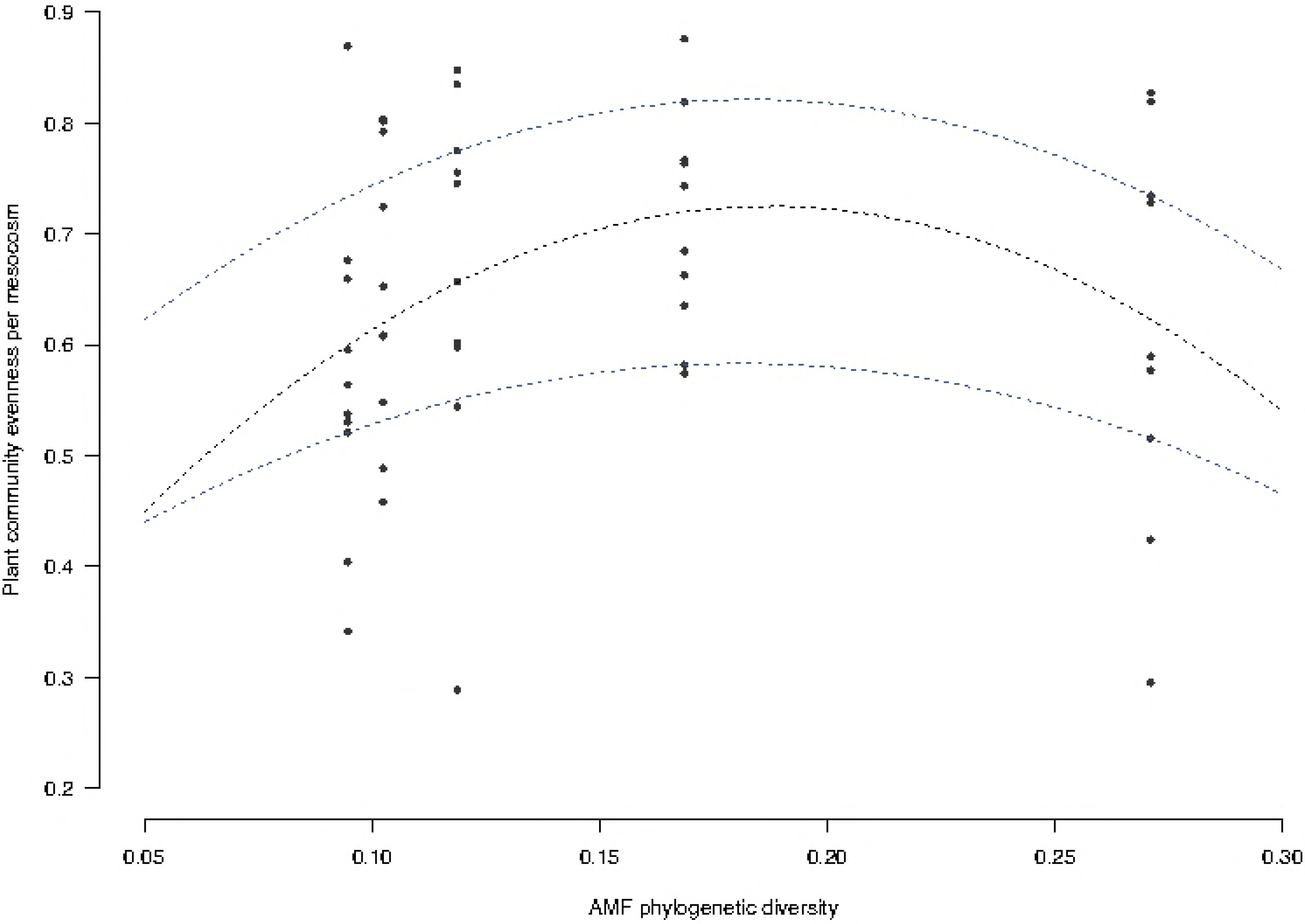
Relationship between plant community evenness and AMF phylogenetic diversity of the five treatments (10-14) with mixed AMF inoculant.

## Discussion

Up to now, the interaction between AMF and plant species within a community was mostly investigated by manipulating AMF at the species level, rarely taking into account the phylogenetic relatedness of AMF isolates. To our knowledge, this study is the first to consider genetic differences among isolates of an AMF species as a source of potential functional trait variability that could differentially impact plant-plant interactions within a community. We show that despite the strong AMF intraspecific effect on single plant responses observed elsewhere [15–16, 18], the effect of *R. irregularis* isolates on each plant species and on the global community were weak. Though weak, the level of *R. irregularis* isolate relatedness did influence the response of some plant species in the community. Furthermore, we observed a phylogenetic conservatism on plant community metrics of responsiveness and on community evenness.

### i) Do different isolates of R. irregularis, or combinations of the isolates, differentially colonize plants and differentially affect plant responsiveness, community structure and productivity?

Although we observed significant differences in AMF colonization ability among the AMF isolates the overall mycorrhizal responsiveness of the community to these isolates did not differ significantly. This suggests that none of the *R. irregularis* strains used in this study was a general plant community growth promoter. Although the mycorrhizal responsiveness of the 6 plant species differed greatly, this was not influenced by the identity of the AMF isolates and, thus, there were no strongly observable differences in plant community structure and productivity due to the different AMF isolate treatments. Despite that number of previously published studies have shown differential effects of within-species AMF variation on the growth of different plant species, in single plant - single fungus experiments, our study did not support the hypothesis that such effects could lead to strong differences at the plant community level. Some of the plant species and AMF isolates used in this study had been shown in previous studies to differentially alter the growth of different plant species.

We found strong positive responsiveness of the subordinate plants of the community except in *L. corniculatus*. The two dominants plants, the grasses *F. pratensis* and *A. eliatus* responded negatively to being colonized by AMF. This resulted in fairly similar mesocosm productivity in every AMF treatment. This observation might be explained by a limited amount of resources available in mesocosms leading to a saturation of the productivity. Such saturation has already been observed in similar experiments [3]. Therefore, the most relevant variation on plant community structure detectable in such an experimental design is the relative contribution of individual plants to community productivity. This is given by the evenness index, which describes the relative competition strength within the plant community [40]. Here, we did not find strong differences in evenness among treatments with different isolates. However, an increase in plant community evenness was mainly due to a decrease in *F. pratensis* drymass relative to an increase in *T. pratense* and *A.elatius*, as these three species contributed to almost 80% of the mesocosm productivity. This effect of dominance mediation by AMF has previously been observed in calcareous grasslands [10].

### ii) Do individual plant species of a community respond more similarly to more closely genetically related AMF isolates compared to more distant ones?

When data collected for each plant species was analysed separately, a phylogenetic signal was detected on AMF colonization of *F. pratensis* meaning that the compatibility of *F. pratensis* and *R. irregularis* in a plant community is linked to the evolutionary history of the different isolates of *R. irregularis*. In addition, a significant phylogenetic signal in mycorrhizal responsiveness of *F. pratensis* as well as for *K. arvensis* was found. This suggests that the trait evolution of different *R. irregularis* isolates not only impacts the ability of the fungus to colonize a given plant species but also indicates that the outcome of the symbiosis in terms of plant growth is more likely to be similar of the AMF isolates are genetically closer. The potential preference of some *R. irregularis* genotypes for some plant species was previously observed [41] as similar *R. irregularis* genotypes tended to be more likely isolated from one plant species that was used as trap-culture than another plant species. This heterogeneity of preference among the plants could be explained by a slight co-adaptations between AMF genotypes and plant species and might favour diversity of AMF and allow the coexistence of closely related AMF genotypes within a community.

### iii) Is the mycorrhizal responsiveness and structure of a plant community more similar in response to more closely genetically related AMF isolates compared to more distant isolates?

At the community level, mean mycorrhizal responsiveness of the community, as well as mean community evenness, were found to be significantly affected by the phylogenetic relationships of the *R. irregularis* isolates. These findings are coherent with the AMF phylogenetic effects observed on *F. pratensis* since community evenness was mainly associated with a decrease of relative ADM contribution of this plant species. *F. pratensis* was the strongly dominant plant of the mesocosm and, similarly to other grass species, *F. pratensis* is known to produce a large amount of roots compared to forbs or legumes [27]. Thus, we could expect that isolates of Gp4 would benefit more than other *R. irregularis* genotypes on dominant grasses. In contrast, members of Gp3 would more likely be associated with increased benefits on subordinate plants since this clade provoked a strong decrease in *F. pratensis* responsiveness, and concomitantly, an increase in *K.arvensis* and *T. pratense* responsiveness. The differences found among these phylogenetic groups suggest that some functional differences might exist at the intra-specific level in *R. irregularis*.

Increasing fungal taxonomic diversity by inoculating different mixes of several *R. irregularis* isolates on the same mesocosm did not significantly influence plant community productivity. Nevertheless, studies showing this relationship were mainly performed in field assays, where space and soil nutrient levels are less limited and, thus, saturation is not reached [3]. We found a significant quadratic relationship between AMF community phylogenetic diversity and plant community evenness, even though only five treatments in the experiment were effectively manipulating AMF phylogenetic diversity. Community evenness was the highest at intermediate level of AMF community phylogenetic diversity and dropped to a lower value where all isolates were combined. This observation suggests a saturating effect of the AMF community where the potential benefit of AMF on biomass partitioning within the plant community is no longer observable. Nevertheless, we did not test all the combinations of AMF isolates so the relationship observed might still be attributed to a complementarity effect occurring by chance or to the effect of one particularly good isolate included in the co-inoculations [42]. However, evidence for coexistence of Gp3 and Gp4 in the same grassland has been assessed, but not for the other combinations of phylogroups [23] This favours the hypothesis of less niche overlap between these two phylogroups, as suggested by the phylogenetic signal, which therefore might increase functional diversity in this particular treatment [21].

## Perspectives and Conclusions

To our knowledge, these are the first results to show that within species diversity of these fungi and in particular phylogenetic relatedness can impact mycorrhizal responsiveness of dominant plants in a community and, consequently, biomass partitioning among a community of plants. These results extend the findings of Powell et al. (2009) [14] in that phylogenetic conservatism of AMF functional traits on plant communities can exist within an AMF species and not only between major AMF clades. If confirmed, this feature is interesting because it suggests that the outcomes of plant-fungal and community-fungal interactions are genetically based and could be conserved over evolutionary time. Further studies should focus on AMF traits that are known to be of major importance to plant growth such as level of nutrient acquisition and transfer to the host and consider them in a phylogenetic context at AMF intra-specific level.

## Authors’ contributions

Conceived and designed the experiments: RS and LV. Performed the experiments: RS and LV Analyzed the data: RS and LV. Wrote the paper: RS, LV and IRS.

## Acknowledgements

We thank Nicolas Ruch, Jérôme Wassef, Cynthia Meizoso, Noémie Gambino for their help in harvesting and Nicolo Tartini and Rafael Joss for their help in root staining. We also thank Aleš Látr and Miroslav Vosatka at Symbiom and Yolande Dalpé at GINCO for providing us with AMF isolates. Bioinformatics computations for the phylogenetic tree were performed at the Vital-IT (http://www.vital-it.ch) Center for High Performance Computing of the Swiss Institute of Bioinformatics. This study was funded by the Swiss National Science Foundation (Grant number: 31003A_162549 to IRS).

## Data accessibility

All data are available on request from the corresponding author.

